# Primary-Level Meta-Analysis of Diversity Outbred Mice Identifies a Fasting Plasma Trimethylamine N-Oxide (TMAO) Locus Modified by Sex and Diet

**DOI:** 10.64898/2026.06.19.733321

**Authors:** Kristen J. Sutton, Levi W. Evans, Erik R. Gertz, Dawson Budke, Nazmul Huda, Phoebe Yam, Myungsuk Kim, Jennifer Rutkowsky, Diana Shih, Jaana Hartiala, Daniel Pomp, Aldons J. Lusis, Hooman Allayee, Brian J. Bennett

## Abstract

Trimethylamine n-oxide (TMAO) is a plasma metabolite linked to adverse cardiometabolic health with complex regulation involving diet, sex, and host genetics. We explored the role of these factors in the genetic regulation of TMAO by performing a primary-level meta-analysis in 1,482 female and male Diversity Outbred (DO) mice from five distinct studies conducted in various regions of the United States. We identified a quantitative trait locus (QTL) associated with TMAO concentration at ∼86 megabase pairs on mouse chromosome 12 with a highly significant LOD score of 67.67. Alleles at the chromosome 12 QTL inherited from the Cast/EiJ (CAST) and PWK/PhJ (PWK) mouse strains primarily drove the association with reduced TMAO concentrations. The chromosome 12 QTL remained significant in sex-stratified analyses and the mode of inheritance appeared additive; furthermore, the QTL was regulated by sex-by-genotype and sex-by-diet interactions. Using a CAST/EiJ X C57BL/6J F2 cross, positional candidates were prioritized by eQTL analysis. Further analysis in a study utilizing the eight DO founding strains identified that *Acyp1* was differentially expressed in hepatic tissue from CAST mice, prompting investigation into its genetic regulation. *Acyp1* demonstrated relevant *cis*- and *trans*-regulation and was significantly correlated with TMAO and hepatic *Fmo3.* However, no significant relationships between *Acyp1* and TMAO were identified in mice inactivated for *Acyp1* or with AAV overexpression of *Acyp1* in the liver. Genes within the chromosome 12 QTL have synteny with humans and may translate to the genetic regulation of human plasma TMAO concentrations and atherosclerosis.

**Author Summary:** We explored the roles of diet, sex, and genetics on the regulation of fasting plasma trimethylamine n-oxide (TMAO) concentration by performing a meta-analysis in 1,482 female and male Diversity Outbred (DO) mice from five unique studies. We identified a QTL associated with TMAO concentration on chromosome 12 at ∼86 mega base pair (Mb) with a highly significant LOD score of 67.67. The locus is modified by both sex and diet.

## Introduction

Trimethylamine n-oxide (TMAO) is a diet-microbe derived metabolite demonstrated to disrupt glucose signaling and cholesterol transport and to promote atherothrombosis and inflammatory pathways (1–5). The relationship between elevated fasting plasma TMAO and cardiometabolic disease is consistent across type 2 diabetes, hypertension, chronic kidney disease, and cardiovascular diseases (6–8). Due to the prevalence of these diseases, reducing TMAO concentration may have beneficial health effects.

Circulating trimethylamine (TMA) is oxidized in the liver by flavin monooxygenase 3 (Fmo3), making the enzyme a candidate for influencing TMAO concentration. Several common SNPs have been characterized in the exomes of the *Fmo3* gene which may impact Fmo3 metabolism and TMAO concentration. In humans, severe decreases in FMO3 activity due to genetic variants are characterized by increased plasma TMA levels and trimethylaminuria disease, which is exacerbated during menstruation in affected women (9) or after consuming foods with TMAO or its precursors choline and carnitine (10). In line with this, the *FMO3* gene has an estrogen response element in its promoter, demonstrating how the genetic architecture may regulate TMAO concentration (11). In mice, *Fmo3* expression is up regulated by estrogen and down regulated by testosterone, contributing to the significantly higher TMAO concentrations observed in female versus male mice (12).

Other studies have addressed the genetic regulation of TMAO using unbiased screening approaches. The Hybrid Mouse Diversity Panel, which includes over ∼100 inbred strains identified a locus for TMAO concentration on chromosome 3 that colocalized with a cis-expression quantitative trait locus (QTL) for the zinc transporter solute carrier family 30 member 7 (*Slc30a7*, *Znt7*) protein (13). Our previous genome-wide association study (GWAS) in DO female mice identified a >3 mega base pair (Mb) wide QTL on chromosome 12 associated with plasma concentrations of TMAO (14). Studies in humans also suggest that the regulation of plasma TMAO is influenced by genetics. A study utilizing parents and children found concordance between the dyads (effect size = 37%) (15). In humans, a GWAS including 1,973 individuals identified two suggestive SNPs; however, neither proved robust in validation studies (13). Subsequent studies in cohorts less than 1,500 people have found few genome-wide and epigenome-wide significant hits related to TMAO (16,17). It is likely that previous attempts to discern the genetic regulation of TMAO in humans were underpowered.

To identify effective lifestyle strategies and potential therapeutic targets that reduce TMAO levels, the roles of diet and genetics require more precise definitions. In the present study, we leveraged 1,482 DO mice to conduct a primary-level meta-analysis examining the genetic regulation of fasting plasma TMAO concentration. Subsequently, we conducted a parallel arm feeding study to investigate the effect of diet on the genetic regulation of TMAO, hypothesizing that a diet enriched in choline, a precursor of TMAO, would elicit the genetic machinery driving plasma TMAO concentration. Utilizing RNA expression analyses from multiple mouse models we identify candidate genes associated with TMAO levels and carry out validation experiments to test their significance. Our study highlights a novel therapeutic target to control plasma TMAO concentration and has implications for human cardiometabolic disease.

## Results

### Description of the Five DO Studies

To investigate the genetic regulation of fasting plasma TMAO concentration, we conducted a primary-level meta-analysis including 1,482 mice (male=514, female=968). The meta-analysis consisted of five independent studies conducted in different regions of the United States, at three different vivaria with varying housing conditions over a decade of time (**Table 1**, study designs in **Figure S1**). Of the studies, four utilized DO mice (n=1028) and one utilized DO-F1 mice (n=454, **methods**). Mice ranged in age from 6 to 41 weeks of age and spanned DO generations 10-44. In all the studies, mice were provided a purified AIN-76A diet at the time-point studied. The mean plasma TMAO concentration on the AIN-76A diet was 2.11 ± 3.10 µM but ranged from 0.11 to 266.12 µM among all mice and was significantly higher in females than males (2.60 ± 2.99 µM versus 1.42 ± 2.97 µM; P<0.001). Together, the primary-level data available for a meta-analysis provided unprecedented power to assess the genetic regulation of plasma TMAO concentration using a systems genetics strategy with mice in a consistent diet environment.

**Table 1.** Descriptions of the five studies in the primary level meta-analysis evaluating the genetic regulation of TMAO including a total of 1,482 Diversity Outbred mice. Mice included in the meta-analysis were housed in different vivarium, ranged in age from 6 to 44 weeks, and were from DO generations 10-44.

| Study Name | N (F) | Gen. | TMAO ($\mu\text{M}$) | | | Age (Wks) | Diet | City, State (Vivarium) |
| --- | --- | --- | --- | --- | --- | --- | --- | --- |
|  |  |  | All Mice | Females | Males |  |  |  |
| DO-UNC1 | 193 (87) | 10 | 10.6 ± 27.5 | 15.93 ± 33.88 | 6.38 ± 20.05 | 8 | AIN-76A | Chapel Hill, NC (Genetic Medicine Building) |
| DO-UNC2 | 261 (261) | 11 | 1.25 ± 0.91 | 1.25 ± 0.91 | NA | 6 | AIN-76A | Chapel Hill, NC (Genetic Medicine Building) |
| DO-Davis | 190 (190) | 26, 28 | 4.07 ± 3.67 | 4.07 ± 3.67 | NA | 40-44 | AIN-76A | Davis, CA (Mouse Biology Program) |
| DO-F1* | 454 (232) | 26, 28 | 6.06 ± 7.31 | 8.88 ± 9.18 | 3.13 ± 2.13 | 8 | AIN-76A | Davis, CA (Mouse Biology Program) |
| DO-Choline | 384 (198) | 42-44 | 1.58 ± 1.52 | 2.20 ± 1.75 | 0.94 ± 0.84 | 7-9 | AIN-76A | Davis, CA (Meyer Hall) |

\*Mice from the DO-F1 study are the F1 progeny of DO-Davis and CETP/ApoE3 Leiden mice. N, Number; F, Female; Gen., Generation.

### Fasting Plasma TMAO Concentration is Genetically Regulated

We first assessed the narrow sense (additive) heritability of plasma TMAO concentration using all 1,482 mice. The heritability of TMAO averaged 53.8%. Heritability scores for each study ranged from 41% to 66%, indicating genetics explains a considerable proportion of TMAO variability in the controlled environment (**Table 2**).

**Table 2.**
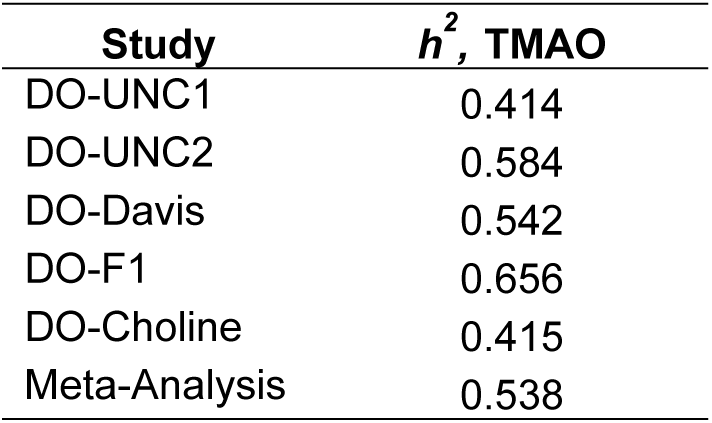
Narrow sense (additive) heritability scores of fasting TMAO concentration from each study in the primary level meta-analysis. Heritability values demonstrate that genetics explains a considerable amount of the metabolite’s variability.

Next, we assessed the genetic regulation of plasma TMAO by performing a genome scan in all mice while controlling for sex, experiment, and kinship. We observed a stark QTL associated with TMAO on chromosome 12 that yielded a LOD score of 67.67 (**Figure 1A**). Notably, this association signal was observed in each of the DO cohorts when evaluated independently and became highly significant in the meta-analysis (**Figure 1B-F**). The 95% Bayesian credible interval was 0.31 Mb (85.68-85.99 Mb) with a peak marker occurring at 85.88 Mb within the gene tubulin tyrosine ligase like 5 (*Ttll5*), which is syntenic to human chromosome 14q24.3. An expanded interval including 1 Mb up and downstream of the peak marker is provided in **Table S2**. Despite our large sample size (n=1,482), the only genome-wide significant QTL associated with fasting plasma TMAO concentration was the QTL on chromosome 12.

**Figure 1.**
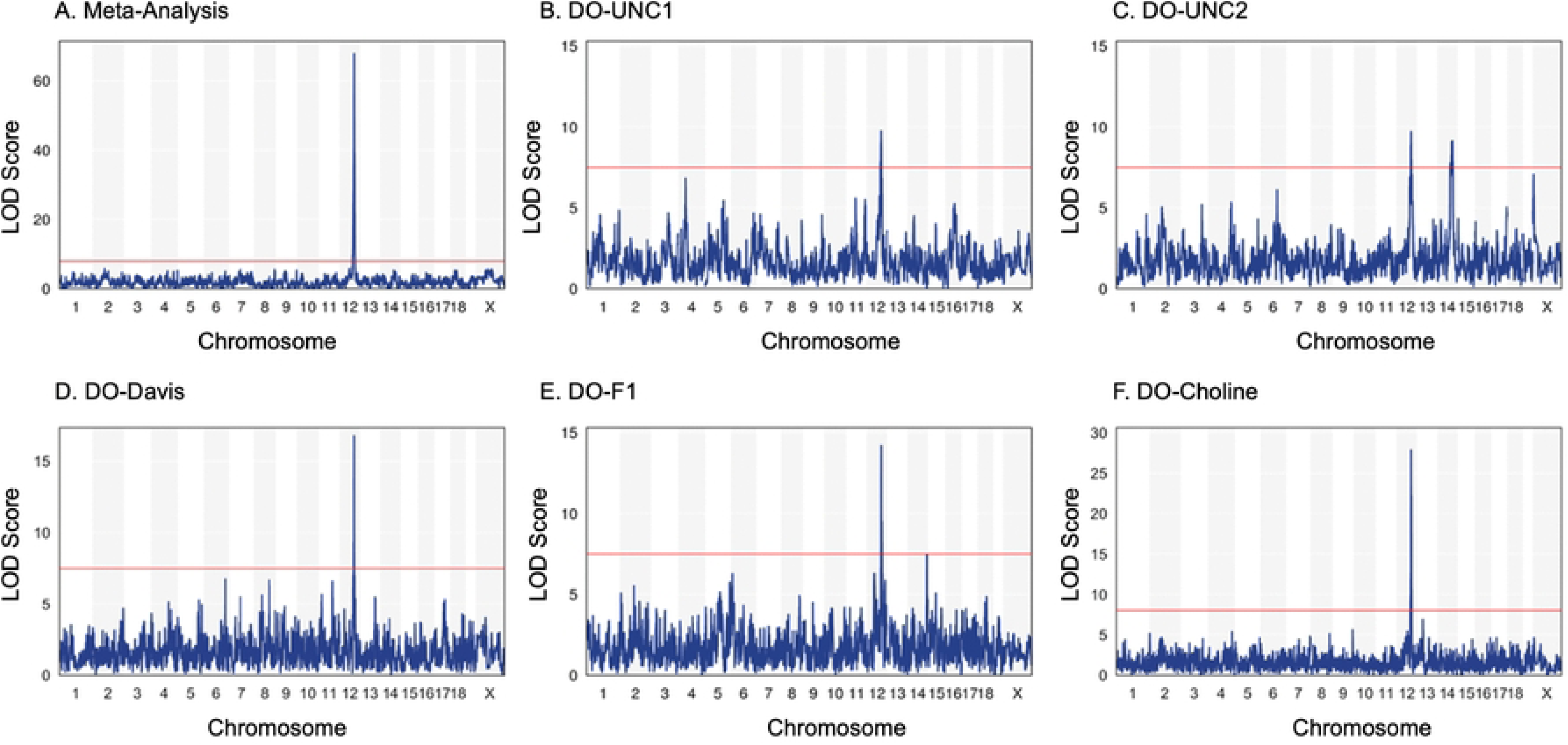
Manhattan plots demonstrating the genome wide regulation of plasma TMAO concentration. Manhattan plots evaluating the genetic regulation of fasting plasma TMAO concentration in the meta-analysis (A) and each of the five studies independently (B-F). A clear QTL on chromosome 12 at ∼86 Mb is identified in each study. The red horizontal lines occur at an LOD score of 7.5.

To further investigate the chromosome 12 QTL, we performed a conditional analysis including the 21 genotyped markers in the 95% Bayesian credible interval. Adding the peak marker to the genome scan completely abolished the significant QTL (**Figure 2A**). The degree to which each marker reduced the LOD score is given in **Table S3**. Ten markers reduced the LOD score below the significance threshold of 7.55, while ten other markers reduced the LOD score, but the QTL remained significant, and 1 marker had no effect on the LOD score. These observations led us to hypothesize that SNPs inherited from one strain may have different effects than those inherited by other founder strains and directed us to investigate the effects of allelotype at the locus. To begin, we leveraged a linear regression model assessing the relationship between TMAO concentration and the probability that an allele was inherited from a given strain. At the marker with the highest LOD score, regression coefficients demonstrate that the QTL is driven by the CAST and PWK strains in each of the five cohorts in the meta-analysis. Further, mice that inherited CAST and/or PWK alleles exhibited reduced fasting plasma TMAO concentration (**Figure 2B**).

**Figure 2.**
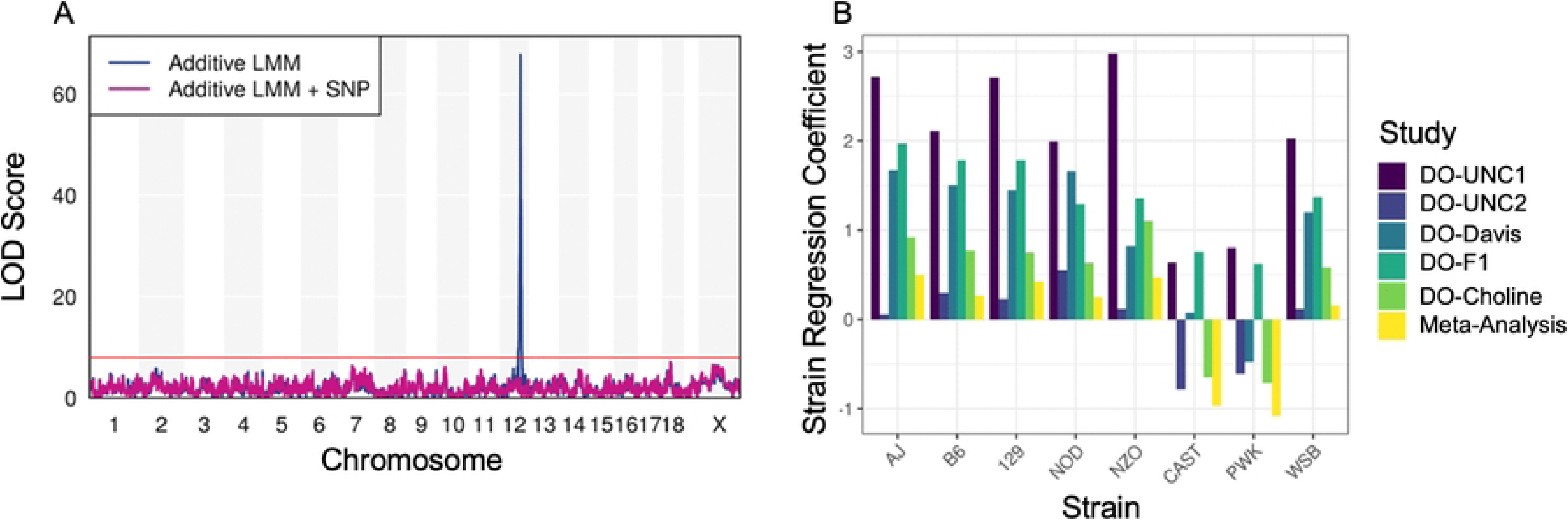
SNP and strain effects at the chromosome 12 QTL. Interrogating the SNP effects at the chromosome 12 QTL associated with fasting plasma TMAO concentration. (A) Adding the SNP with the highest LOD score (peak SNP) as an additive covariate in the linear mixed model eliminates the significant QTL on chromosome 12. (B) At the peak SNP, alleles inherited from CAST and PWK mice are inversely associated with TMAO concentrations across each study in the meta-analysis. Red horizontal line = LOD score of 7.5.

### Role of Sex in the Genetic Regulation of Plasma TMAO Concentration

Next, we explored whether the genetic regulation of plasma TMAO concentration was sex specific by repeating the meta-analysis using sex-stratified genome scans while controlling for experiment and relatedness. The QTL on chromosome 12 was replicated in both sexes and remained highly significant in female (LOD=43.8) and male mice (LOD=24.4) (**Figure 3A magenta** and **green)**. Furthermore, TMAO concentrations as a function of genotype at the peak marker revealed a sex by genotype interaction (P=0.0487) (**Figure 3B**), where male mice homozygous for the effect allele had significantly greater reductions in TMAO concentration versus female mice (β=-0.681, P<0.001). Although the mode of inheritance appeared additive in both sexes, the effect of inheriting two variant alleles in males had a stronger reduction in TMAO concentration than females of the same genotype. The gene-by-sex interaction also replicated when these analyses were conducted independently in the DO-Choline study (P=0.014) (**Figure S2**).

**Figure 3.**
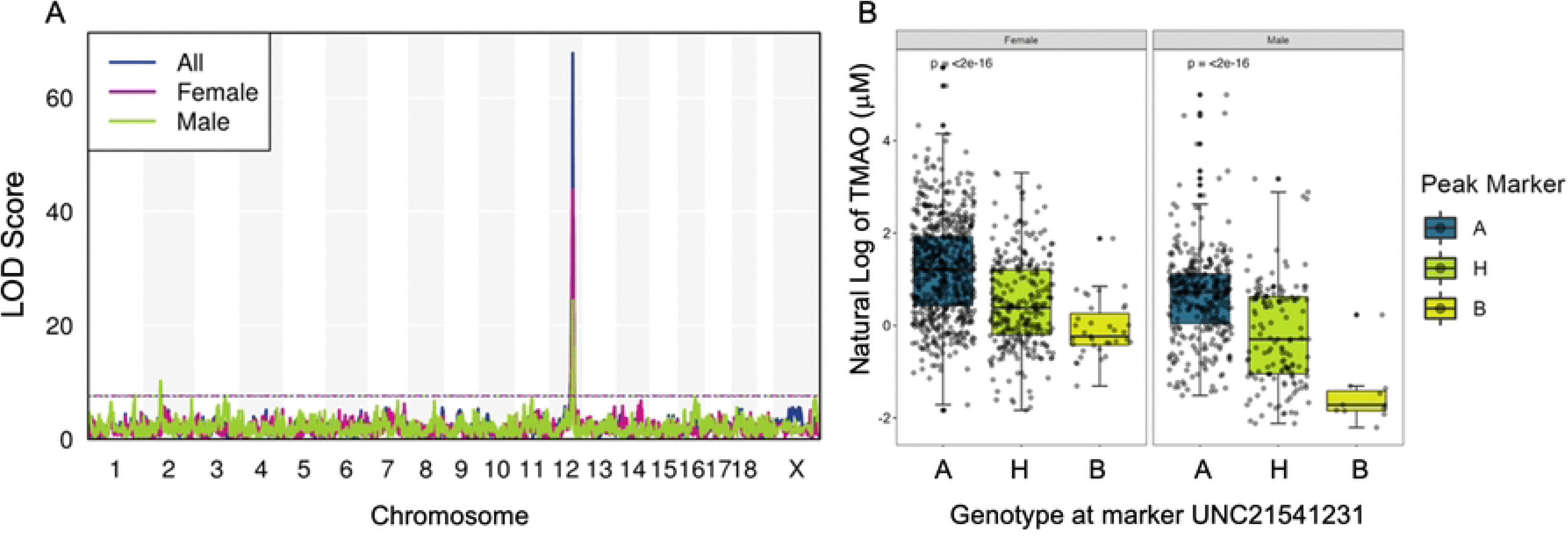
Chromosome 12 QTL is affected by sex. The QTL on chromosome 12 associated with fasting plasma TMAO concentration is affected by sex. (A) Manhattan plot showing the genome wide results of the linear mixed model assessing plasma TMAO concentration in all mice (purple), female mice (magenta), and male mice (lime green). A narrow interval is identified in all three mouse subsets as indicated by the sharp vertical line on chromosome 12. (B) Distribution of TMAO concentration by genotype and sex at the peak SNP within the QTL on chromosome 12 associated with fasting plasma TMAO concentration. ANOVA was used to determine differences in means of TMAO concentration by genotype and corresponds to the printed P-values.

The difference in LOD scores between the sexes may be due to differences in sample size or due to other sex-specific genetic factors that influence the complex trait. In female mice, no other TMAO-associating QTL were identified. However, in male mice, three additional QTL surpassed the 95% threshold of 7.56 determined by 1000 permutations (**Table 3**). The first locus was located on chromosome 1 at 165.49 Mb (95% BCI, 81.29-165.56 Mb) and a LOD score of 7.71. The confidence interval for this QTL was large (84.96 Mb wide) but notably contained the genes *Fmo1*, *Fmo2*, *Fmo3*, and *Fmo4* located between 162.62 and 162.81 Mb (**Table S4**). The second locus was located on chromosome 2 at 63.87 Mb and a LOD score of 10.21. The interval for this QTL was smaller (3.34 Mb) but did not contain an obvious candidate gene (**Table S5**). The third locus was located on chromosome 3 at 110.59 Mb (87.13-123.52 Mb) with a LOD score of 7.65. Although broad (36.28 Mb wide) the interval contained *Fmo5* and *Znt7* (**Table S6**), genes previously linked to TMAO concentration.

**Table 3.** Genetic regulation of TMAO in male and female mice, separately.

| Mice | Phenotype | chromosome | Position (Mb) | LOD | 95% BCI (Mb) |
| --- | --- | --- | --- | --- | --- |
| Males | TMAO | 1 | 165.492 | 7.71 | 81.291-165.564 |
|  |  | 2 | 63.872 | 10.21 | 62.526-66.004 |
|  |  | 3 | 110.591 | 7.65 | 87.131-123.520 |
|  |  | 12 | 85.862 | 24.4 | 85.677-86.081 |
| Females | TMAO | 12 | 85.876 | 43.83 | 85.677-86.034 |

chromosome, chromosome; Mb, mega base pair; LOD, logarithm of odds; BCI, Bayesian credible interval.

### TMAO Concentration is Impacted by Dietary Choline

In addition to the role of genetics, plasma TMAO concentration is dependent on the dietary intake of TMAO precursors including choline and L-carnitine. Therefore, we next investigated whether the genetic regulation of TMAO was impacted by administration of a high-choline (HC) diet using the DO-Choline study. We hypothesized that the HC diet would elicit the genetic machinery required to generate, process, or eliminate TMAO, thus providing an opportunity to observe gene-by-diet interactions.

At baseline, mice on the AIN-76A diet had mean ± standard deviation plasma TMAO concentrations of 1.56 ± 1.50 µM, which significantly increased (P<0.001) to 25.96 ± 43.27 µM in mice randomized to the HC diet. The distributions of TMAO concentrations by diet and sex are shown in **Figure 4A**. Regardless of the diet, female mice had higher plasma TMAO concentration than male mice. At baseline when all mice were consuming AIN-76A, the average TMAO concentration was about 2 times higher in females than males (female, 2.20 ± 1.75 µM; male, 0.94 ± 0.83 µM; P<0.001). In comparison, mice randomized to the HC diet had dramatic but disparate increases in TMAO; female mice had approximately 3 times higher TMAO concentration than male mice (female, 38.43 ± 57.11 µM; male: 12.91 ± 10.49 µM; P < 0.001). We then compared baseline to final TMAO concentrations. Male mice that always consumed AIN-76A had a Spearman correlation value of 0.71 compared to 0.67 in females (**Figure 4B**), while male mice who switched to the HC diet had a correlation value of 0.29 versus 0.18 in females (**Figure 4C**). For mice who switched diets, there was a non-significant difference between the correlations by sex (P=0.36). Overall, these results suggest that within a diet treatment mice who had high concentrations at baseline tended to have higher values at follow-up, but that diet can dramatically affect TMAO values.

**Figure 4.**
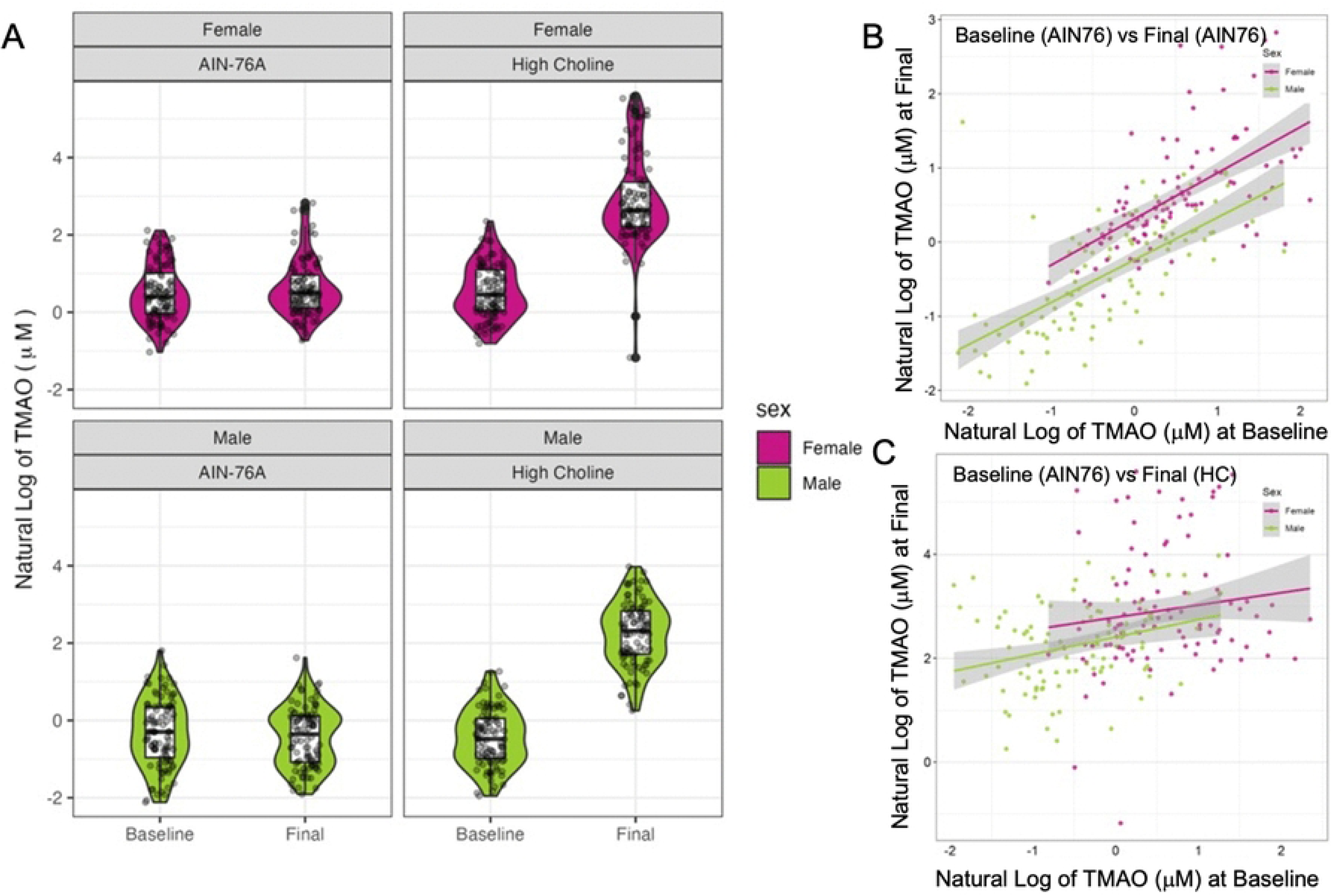
TMAO concentrations across sex and diet. Distribution of fasting plasma TMAO concentrations in female and male DO mice in different diet conditions from the DO-Choline study. (A) Violin plots demonstrate the effects of sex and diet on TMAO. The plots in the “AIN-76A” column (left) represent mice who were randomized to the AIN76 study arm while plots in the “High Choline” column (right) represent mice who were randomized to the HC diet arm. (B) Correlation of TMAO concentrations at baseline and final in mice who consumed AIN76 throughout the duration of the study. (C) Correlation of TMAO at baseline and final in mice who were randomized to the HC diet arm. Correlations were conducted via the Spearman correlation test.

We then assessed the genetic regulation of TMAO under different dietary conditions and observed that the chromosome 12 QTL was diet-specific (**Figure 5A**). For mice that continued to be fed the AIN-76A diet, the Chr 12 QTL was replicated with a LOD of 10.7. The 95% BCI interval was only slightly larger at 0.41 Mbp despite the ∼4-fold smaller sample size (**Table S7**). However, no TMAO-associated QTLs were detected following two weeks of the diet enriched for choline (HC) (**Figure 5B**), indicating that the genetic signal derived from the chromosome 12 QTL was diet-specific.

**Figure 5.**
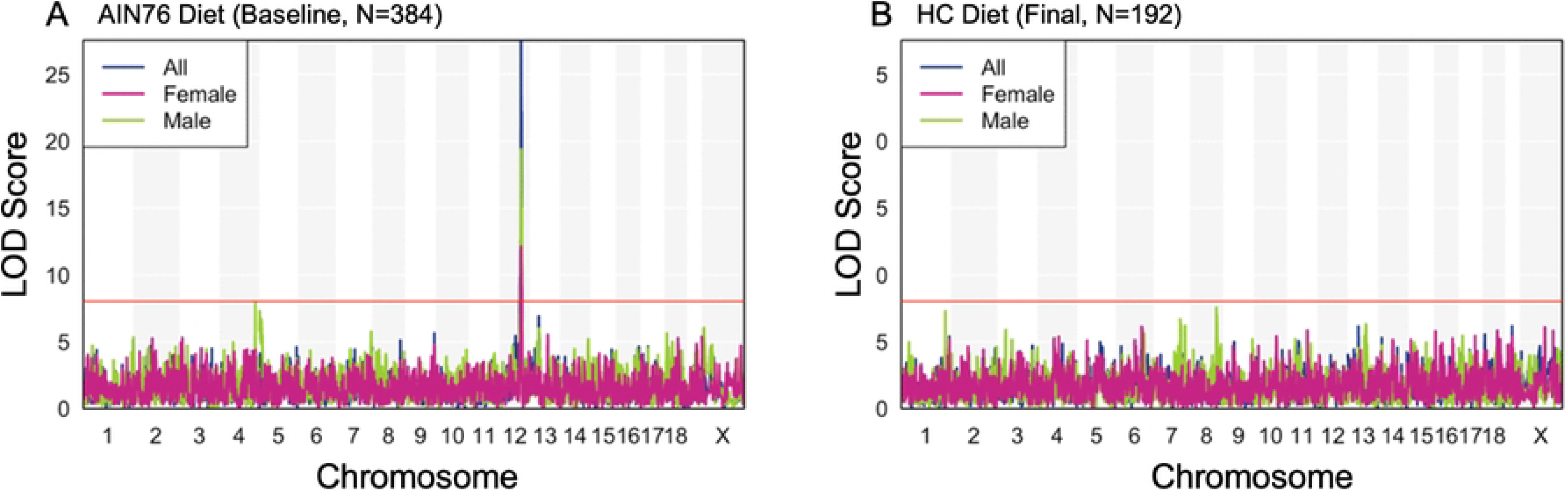
**TMAO is regulated by a gene-by-diet interaction**. (A) Manhattan plot illustrating genome wide LOD scores associated with plasma TMAO concentration in the AIN76 diet environment. A clear QTL is present on Chr 12 with an LOD score of 27.86. The horizontal red line corresponds to an LOD value of 8.03, which signifies the genome wide 95% LOD threshold calculated by conducting 1000 permutations. (B) Similar to A in the HC diet environment. The red line corresponding to the 95% LOD threshold is at 8.24. The QTL on Chr 12 is not observed in the HC diet environment.

### Differential Gene Expression from the Progenitor DO Strains

Next, we caried out a study in the five classically inbred and the three wild-derived inbred strains used to generate the DO (**methods**, DO-Progenitor study) to further interrogate the expression of genes at the TMAO associated locus on chromosome 12. This analysis provided the opportunity to examine the genetic effects of each founding strain and to determine whether the expression of genes within the QTL matched the regression coefficient pattern from the previous analyses. From the regression coefficient pattern observed in **Figure 2B**, we hypothesized that the expression of genes, specifically in the liver, that were similar between CAST and PWK, but different from the other six strains, may play an important role in regulating plasma TMAO concentrations. We used a conservative approach that included 22 genes within a 1 Mb window of the highest-LOD scoring SNP at 86 Mb. We performed differential expression analyses by strain and identified 12 genes whose mRNA levels differed significantly across strains (Kruskal-Wallis, Padj<0.05) (**Figure 6A**). The complete list of differential expression results is provided in **Table S8**. Of these genes, the top four had unique expression patterns between CAST and/or PWK strains versus most other strains, including angel homolog 1 (*Angel1)* **(Figure 6B)**, ribosomal protein S6 kinase like 1 (*Rps6kl1)* (**Figure 6C**), estrogen-related receptor beta *(Esrrb)* (**Figure 6D**), and acylphosphatase 1 erythrocyte common type *(Acyp1)* (**Figure 6E**).

**Figure 6.**
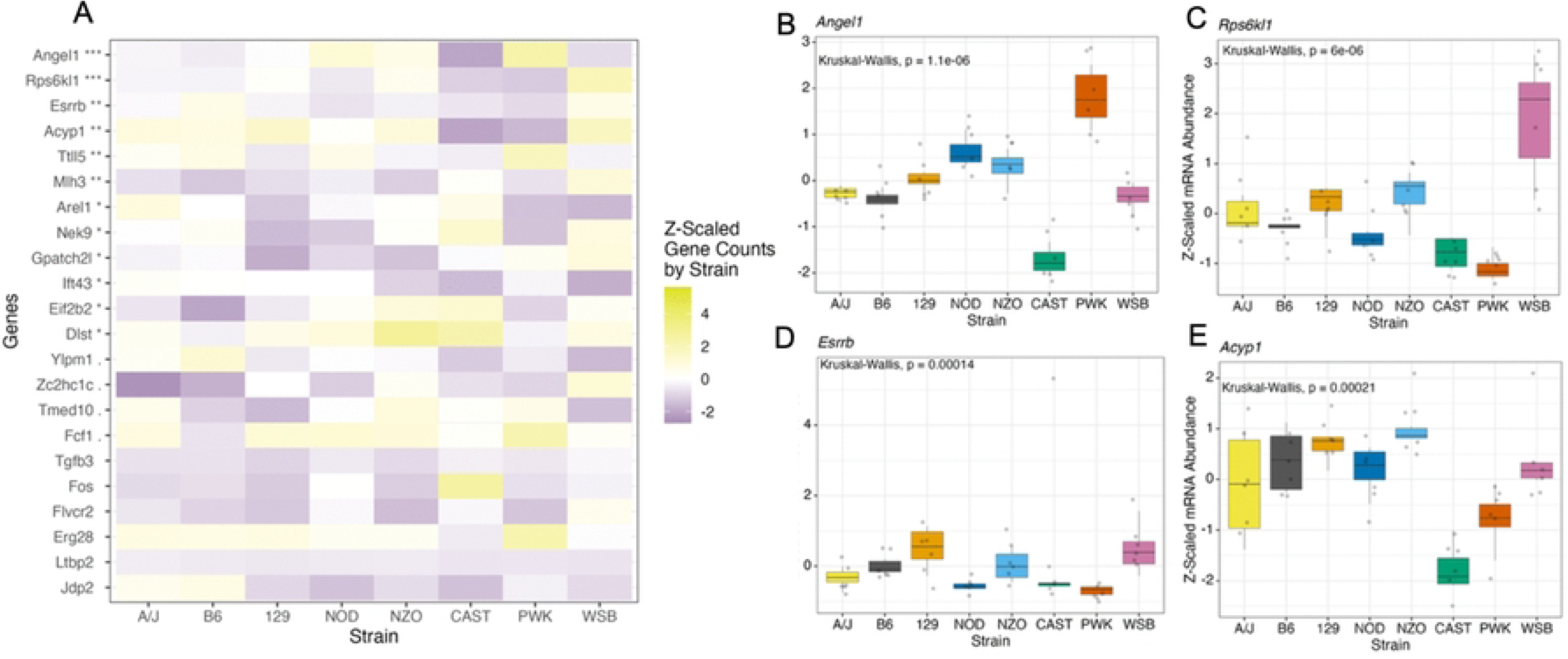
Hepatic expression of candidate genes from the chromosome 12 QTL. Assessing the hepatic expression of candidate genes by strain from the DO-Progenitor study. (A) Differences in hepatic RNA expression of genes within the chromosome 12 QTL among the eight strains used to generate the DO mouse model. Genes that were significantly different by strain are followed by a symbol where ***, P<0.001; **, P<0.01; *, P<0.05; ., P<0.1. (B-E) RNA expression of the top 4 most significant genes is illustrated. (B) *Angel1* shows significantly different expression levels between CAST and PWK strains (P=0.004). (C) CAST has reduced gene expression of *Rps6kl1* versus all strains (P<0.05) but PWK (P=0.08) and NOD (P=0.33). PWK has reduced gene expression of *Rps6kl1* versus all strains (P<0.05) but CAST (P=0.08). (D) PWK has reduced expression of *Esrrb* than B6 (P=0.015), 129 (P=0.015), and NZO (P=0.015). (E) CAST has reduced gene expression of *Acyp1* versus all strains (P<0.05). PWK has reduced gene expression of *Acyp1* versus all strains (P<0.05) but AJ (P=0.5). Statistical tests were completed by the Kruskal Wallis test and the pairwise Wilcox test. Tests were corrected for multiple comparisons using the BH method. Gene counts were Z scaled for graphing purposes.

### Expression-QTL Analyses Support CAST Allele Contribution to Locus

We then leveraged expression data generated from liver, adipose, muscle, and brain tissue derived from a C57BL/6J X CAST/Ei F2 intercross (18). This model enabled us to investigate local regulatory effects that may be influencing the chromosome 12 QTL and TMAO levels, and to assess the role of the CAST strain. In all four tissues, we identified several eQTL (LOD > 7) for positional candidates at the chromosome 12 QTL identified in the meta-analysis. We restricted our assessment to *cis-*eQTL whose transcription start site was within 1 Mb of the peak SNP from the meta-analyses’ chromosome 12 QTL. This resulted in identifying 10 genes whose expression in liver, adipose, muscle, and/or brain was associated with a SNP within our locus (**Table 4**). Among these genes, *Acyp1*, mutL homolog 3 (*Mlh3*), and eukaryotic translation initiation factor 2B, subunit 2 beta (*eIF-2ß2*) had significant eQTL in liver (**Table 3**) and brain (**Table S9**) in analyses with both males and females, as well as in sex-stratified analyses. The *cis-*eQTL identified through these analyses thus provided support that genetic variants harbored by the CAST strain could contribute to the observed QTL associated with fasting plasma TMAO concentration on chromosome 12.

**Table 4.** Expression QTL overlapping chromosome 12 QTL. Expression QTL analysis from liver tissue derived from a CAST X BL6 F2 mouse model identifies several cis-regulated genes within a 1 Mb window of the maximal SNP in the chromosome 12 QTL.

| Sex | chromosome | LOD | eQTL<br>Max<br>Position<br>(CM) | eQTL<br>Max<br>Position<br>(Mb) | Transcript<br>start (Mb) | Transcript<br>Gene<br>Abbreviation |
| --- | --- | --- | --- | --- | --- | --- |
| All | 12 | 104.9 | 36.94 | 86.434 | 85.335 | Acyp1 |
|  |  | 83.0 | 36.94 | 86.434 | 85.317 | Mlh3 |
|  |  | 51.6 | 36.94 | 86.434 | 85.266 | Eif2b2 |
|  |  | 34.9 | 36.94 | 86.434 | 85.158 | Dlst |
|  |  | 26.1 | 37.1 | 86.562 | 85.317 | Mlh3 |
|  |  | 12.6 | 36.94 | 86.434 | 85.421 | Tmed10 |
| Female | 12 | 58.9 | 36.94 | 86.434 | 85.335 | Acyp1 |
|  |  | 49.1 | 38.94 | 86.434 | 85.317 | Mlh3 |
|  |  | 32.0 | 36.94 | 86.434 | 85.266 | Eif2b2 |
|  |  | 19.1 | 36.94 | 86.434 | 85.158 | Dlst |
|  |  | 17.0 | 37.1 | 86.562 | 85.317 | Mlh3 |
|  |  | 9.6 | 36.94 | 86.434 | 85.421 | Tmed10 |
| Male | 12 | 46.4 | 36.94 | 86.434 | 85.335 | Acyp1 |
|  |  | 37.1 | 36.94 | 86.434 | 85.317 | Mlh3 |
|  |  | 19.7 | 36.94 | 86.434 | 85.266 | Eif2b2 |

Transcription start site was taken from the Mouse Genome Database by Jackson Laboratory and is relative to the 5' end. chromosome, chromosome; LOD, logarithm of odds; CM, centimorgan; bp, base pair; Acyp1, acylphosphatase 1, erythrocyte (common) type; Mlh3, mutL homolog 3; Eif2b2, eukaryotic translation initiation factor 2B, subunit 2 beta; Dlst, dihydrolipoamide S-succinyltransferase (E2 component of 2-oxo-glutarate complex); Tmed10, transmembrane p24 trafficking protein 10.

We next isolated hepatic RNA from the DO-Choline study and quantified expression of the four genes with significant eQTL in the liver. Controlling for sex, diet, and experimental batch, we identified *cis-*eQTL (LOD>7.5) for the genes *Acyp1* and *Mlh3* (**Table 5**). *Acyp1*, but not *Mlh3*, appeared to be driven by alleles from the CAST strain matching the pattern observed for the TMAO QTL (**Figure S3**). We hypothesized that these genes may affect TMAO concentrations by interacting with *Fmo3*, the main enzyme that oxidizes TMA to TMAO. Expression of *Acyp1* (r=-0.376, P<0.001), *Dlst* (r=0.293, P<0.001), *Eif2b2* (r=0.276, P<0.001), and *Ttll5* (r=0.274, P<0.001), were significantly associated with *Fmo3* expression (**Table 5**). However, only *Acyp1* (r=-0.115, P=0.028), was significantly correlated with fasting plasma TMAO concentrations (**Table 6**).

**Table 5.**
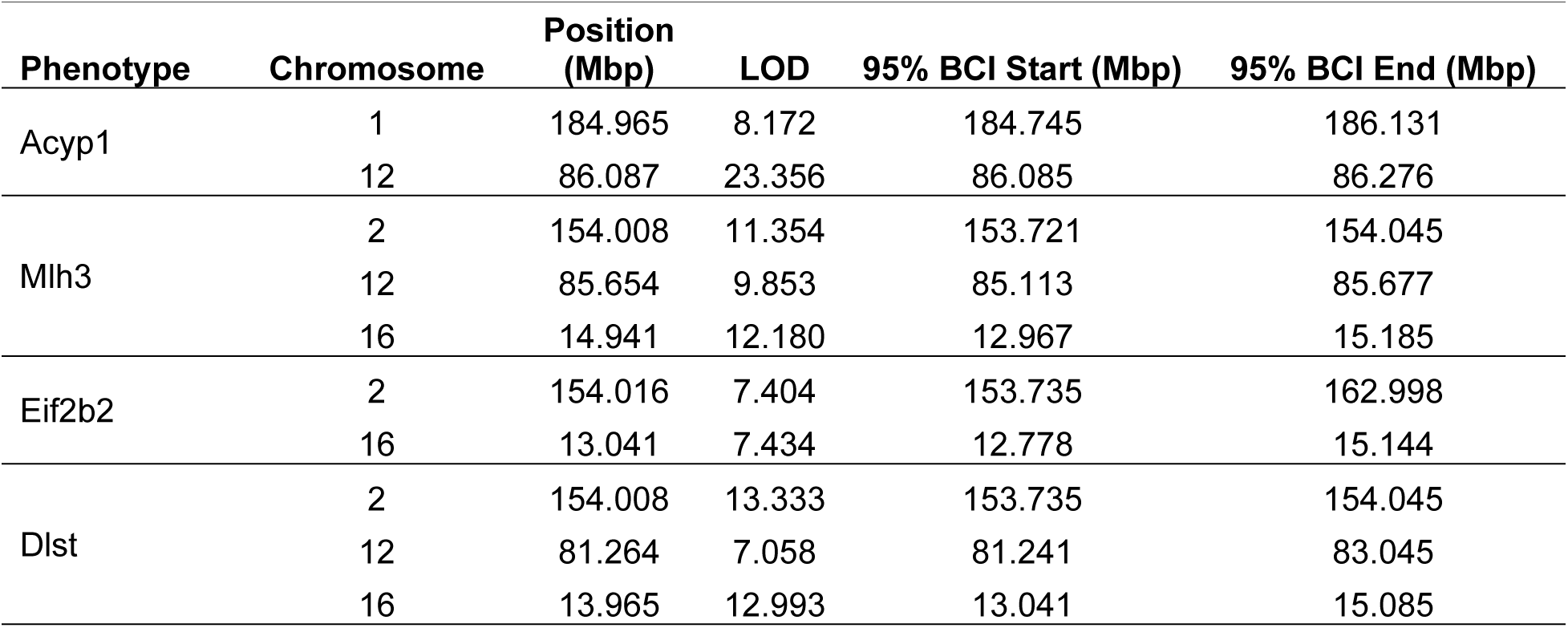
Expression QTL results of candidate genes in the Chr 12 QTL associated with fasting plasma TMAO from the DO-Choline study. Only QTL with an LOD>7 are provided.

| Phenotype | Chromosome | Position (Mbp) | LOD | 95% BCI Start (Mbp) | 95% BCI End (Mbp) |
| --- | --- | --- | --- | --- | --- |
| Acyp1 | 1 | 184.965 | 8.172 | 184.745 | 186.131 |
|  | 12 | 86.087 | 23.356 | 86.085 | 86.276 |
| Mlh3 | 2 | 154.008 | 11.354 | 153.721 | 154.045 |
|  | 12 | 85.654 | 9.853 | 85.113 | 85.677 |
|  | 16 | 14.941 | 12.180 | 12.967 | 15.185 |
| Eif2b2 | 2 | 154.016 | 7.404 | 153.735 | 162.998 |
|  | 16 | 13.041 | 7.434 | 12.778 | 15.144 |
| Dlst | 2 | 154.008 | 13.333 | 153.735 | 154.045 |
|  | 12 | 81.264 | 7.058 | 81.241 | 83.045 |
|  | 16 | 13.965 | 12.993 | 13.041 | 15.085 |

**Table 6.**
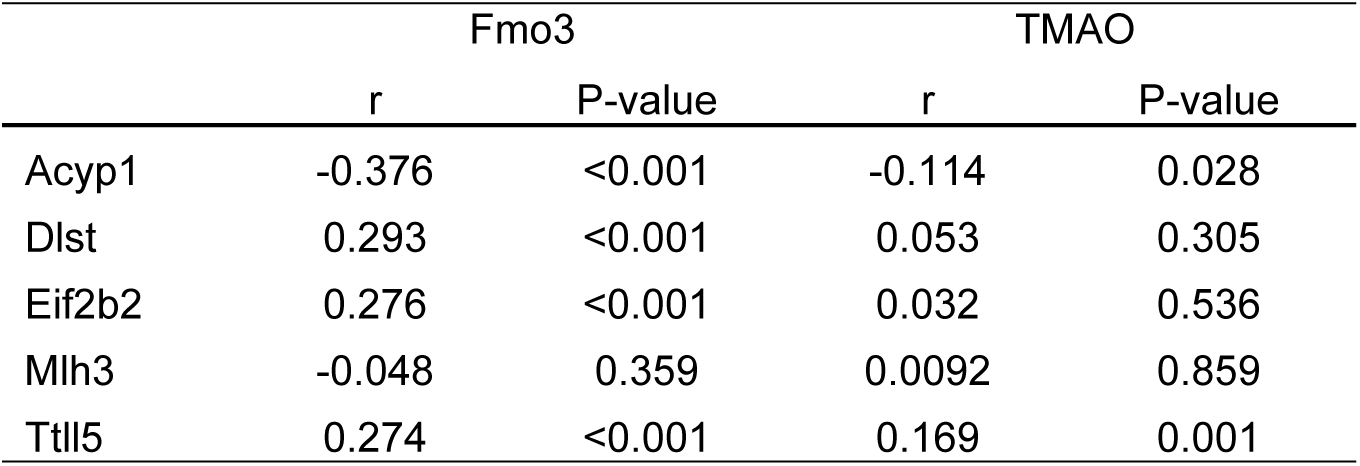
Pearson’s correlations of hepatic expression of candidate genes within the Chr 12 QTL and hepatic FMO expression and plasma TMAO concentrations from the DO-Choline study.

### Candidate Gene Exploration

*Acyp1* had a significant *cis-*eQTL and was correlated to TMAO and *Fmo3* in the DO-Choline and the C57BL/6J X CAST/Ei F2 studies, and expression patterns matching the strain effects in the DO-Progenitor study. Thus, we hypothesized that *Acyp1* could influence TMAO concentrations. We designed under- and over-expression studies utilizing CRISPR and AAV8 technologies, respectively. AAV transduction of hepatic specific *Acyp1* in B6 mice led to approximately a 1.5x upregulation of *Acyp1* with no detectable differences in *Fmo3* or plasma TMAO (**Figure 7 E-H**). Mice with a CRSPR targeted deletion of Acyp1 were rederived and fed AIN-76A diet for 2 weeks. QPCR confirmed a near complete ablation of *Acyp1* expression in the CRISPR mice, but plasma TMAO concentration was also not different from wild-type controls (**Figure 7A-D**)

**Figure 7.**
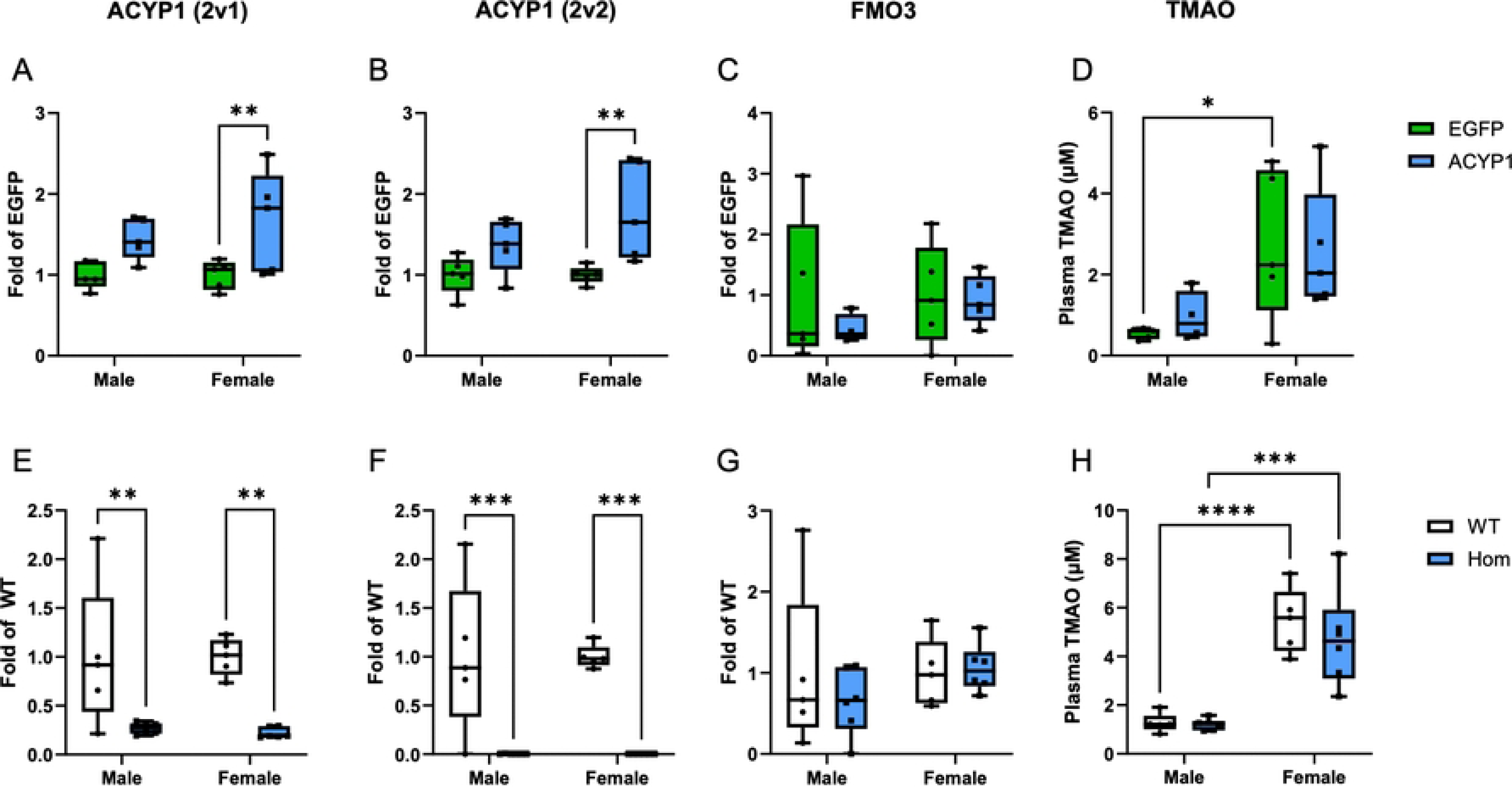
Hepatic expression of Acyp1 and plasma TMAO concentrations in ACYP1 CRISPR knockout mice and in mice post administration of AAV8-ACYP1. (A-C) Fold expression (vs WT) of Acyp1 and Fmo3 in male and female CRISPR knockout mice. (D) Plasma TMAO concentrations were assessed at Baseline and post 2 week feeding of AIN-76A diet. (n=5-6/group). (E-F) Fold expression (vs AAV8-EGFP) of Acyp1 and Fmo3. (H) Plasma concentration of TMAO was assessed in male and female mice 2 weeks post AAV8 injection (n=4-5/group). ****, P<0.0001, ***, P<0.001; **, P<0.01; *, P<0.05.

## Discussion

**Here we employed a primary-level meta-analysis including five unique studies incorporating 1,482 male and female heterogeneous DO mice reared in different vivaria in multiple cites over approximately a decade**. Our studies encompassed multiple life stages including young adulthood to late middle age and included mice who had undergone the stresses and physiological changes associated with pregnancy and birth. Regardless of sex, we identified a highly significant locus on chromosome 12 at 86 Mb that is associated with reduced TMAO concentrations. We performed a series of experiments in complementary mouse models to investigate the strain and gene driving the QTL. We also explored the role of sex at the locus and revealed a sex-by-genotype interaction as well as 3 male-specific QTLs. Furthermore, we investigated the impact of diet on the genetic regulation of plasma TMAO and identified that the HC diet ameliorated the chromosome 12 QTL. Our results provide robust evidence that fasting plasma TMAO concentration is regulated by a QTL on chromosome 12 that is influenced by sex and diet. Moreover, we provide compelling genetic results that may be translatable to humans.

### TMAO Regulation Affected by Sex and Diet

Studies in humans suggest that plasma TMAO has high levels of inter-individual variation (19). Although there is clear genetic regulation at the chromosome 12 locus, diet has a drastic effect on TMAO production that can obscure this regulation. This was demonstrated among the subset of mice that were challenged with a high choline diet in which the QTL for TMAO was not detected despite having 90% power to detect a QTL with an effect size of 5% (20). Although we hypothesized that perturbing the system with high amounts of choline would reveal the genetic machinery regulating TMAO, it is possible that a diet enriched with 1.1% choline saturated the regulatory mechanisms on chromosome 12 involved in TMAO metabolism, such that its ability to impact concentrations was overcome.

The disparate concentrations of plasma TMAO by sex have been previously described in mice (4). This may be partially explained by the sexually dimorphic expression of *Fmo3*, the gene that oxidizes TMA to TMAO (12). In this regard, an estrogen response element has been described upstream of the *Fmo3* transcription site, which could influence *Fmo3* transcription in a sex specific manner (11). Because of these observations, we investigated whether the genetic regulation of plasma TMAO concentration differed by sex. Whether in all mice, or in females and males individually, our analyses identified the significant QTL on chromosome 12 in all cases. In the meta-analysis, the LOD score was highest in the full cohort (LOD 67.8) followed by females (LOD 43.8) and males (LOD 24.4). The lower LOD score in males reflects the reduced number of males in our meta-analysis cohorts (n= 494 out of 1,482 total mice). Additionally, we investigated the relationship between TMAO concentration and the number of inherited effect alleles and consistently observed sex-specific genetic effect sizes. In male mice, inheriting two effect alleles at the peak marker of the chromosome 12 QTL reduced the concentration of plasma TMAO more than in females of the same genotype. Therefore, the homozygous genotype may reduce the limited potential to generate *Fmo3* in males, whereas the homozygous genotype had less effect in females, who begin with greater potential to express *Fmo3*. With relation to humans, a study of ∼360 Caucasians described significantly higher *FMO3* expression in females than males, but the difference in expression was much smaller than in mice (12,21). Thus, the variant on chromosome 12 regulating TMAO concentrations in mice may translate to humans but it is less clear whether the sex-by-genotype interaction will prove robust.

In male mice in the meta-analysis, we identified three additional QTL associated with fasting plasma TMAO concentration. Two of these QTL contained *Fmo* genes 1 through 6 (chromosome 1: *Fmo1*-*Fmo4*, *Fmo6*; chromosome 3: *Fmo5*). Although FMO3 has been shown to oxidize up to ∼90% of TMA, FMO1 and FMO2 have been demonstrated to generate TMAO (12). Thus, it is possible that other *Fmo* genes play more important roles in regulating TMAO concentration in male mice due to their decreased expression of *Fmo3* compared to females. It is unclear how sex specific results in the current DO study relate to the well documented difference in risk of atherosclerosis between younger men and women (22) or the increase in CAD in women who transition into menopausal early (23).

### Candidate Genes Regulating Plasma TMAO Concentration

Although our combined analyses provided approximately 10-times improved genetic resolution from its initial reporting of ∼3 Mbp in (DO-UNC2; n=280, outbreeding generation 11) to 0.31 Mbp (meta-analysis; n=1,482, outbreeding generations 10-44), the approach did not reduce the QTL to a single candidate gene suggesting limits to association mapping techniques due to linkage disequilibrium (LD) and highlighting the remaining challenge of linking the top variant to its putative gene in DO mice. To refine our results to a single candidate gene we used several approaches. First, we examined the roles of strain in the chromosome 12 QTL and observed that the effect allele originated from the CAST strain. Utilizing the DO-Progenitors, many of the transcripts with *cis*-eQTL were differentially expressed in CAST versus the other DO strains, including *Angel1, Rps6kl1, Esrrb, and Acyp1.* The expression of these four genes tended to be differentially expressed in CAST and/or PWK than the other six strains, matching the pattern of the regression coefficients identified in the meta-analysis (**Figure 3**). Using a CAST X BL6 F2 model, we observed several *cis*-eQTL overlapping the chromosome 12 QTL including *Acyp1*. Quantifying hepatic expression of genes in the chromosome 12 locus in the DO-Choline study, we identified significant QTL associated with *Acyp1*, which was correlated with TMAO and *Fmo3* expression. With multiple levels of evidence pointing to *Acyp1* as a leading candidate gene we designed reverse-genetic experiments perturbing *Acyp1* to evaluate its impact on TMAO concentration in mice fed the AIN-76A diet. Despite up and down-regulating hepatic *Acyp1* in two different mouse models, no associations with TMAO were identified. Our candidate selection was based on gene expression and eQTLs, which was one potential mechanism underlying the robust QTL on chromosome 12.

The underlying gene and exact mechanism underlying the TMAO QTL on chromosome 12 remains unknown. We hypothesized that the candidate gene regulates TMAO levels via interaction with *Fmo3* (i.e. generation), but there is limited evidence or protein-protein interactions for the genes in the chromosome 12 locus. Our candidate gene choice relies extensively on mRNA expression in the liver. However, it is possible that *Acyp1* and/or another positional candidate gene at the chromosome 12 QTL influences TMAO concentration in other metabolically relevant tissues, such as the kidney, which is known to excrete TMAO into urine (24). Additionally, protein-protein interactions could be important. For example untargeted proteomics and co-immunoprecipitation have been used to study protein-FMO3 interactions (25). One Chr 12 positional candidate, dihydrolipoamide S-succinyltransferase (*Dlst*), has been shown to interact with FMO3 and in mice given an aryl-hydrocarbon to induce FMO3 expression, co-immunoprecipitation experiments identified *Dlst* as significantly enriched by the FMO3 antibody versus the IgG control. Furthermore, *Dlst* expression increased 1.34-fold in female Fmo3^-/-^ mice on a C57BL/6 background compared to controls, whereas it did not change in female Fmo3^-/-^ mice on an FVB background (25). Additionally, Coffey et al. identified miR-146a-5p was inversely associated with *Fmo3* and had a validated mRNA-miRNA interaction between itself and *Dlst* (14). The same study also observed similar relationships between *Dlst*’s neighboring gene NUMB Endocytic Adaptor Protein (*Numb*) and miR-146a-5p and *Fmo3,* but evidence from our experiments did not support Numb as the likely causal gene at the chromosome 12 QTL.

By leveraging five independent studies conducted in 1,482 mice across different vivaria, at different ages, and in different points of time, we provide compelling genetic evidence that plasma TMAO concentrations are influenced by a locus on mouse chromosome 12. However, despite incorporating multiple follow-up bioinformatic and functional analyses to interrogate the chromosome 12 QTL, the underlying causal gene remains elusive. In all, we only assessed the regulation of plasma TMAO concentration, which may not reflect other important stages of the TMAO pathway, such as TMA generation by the gut microbiome and TMAO excretion via the kidneys. We speculated that the candidate gene influences the expression or activity of the *Fmo3* gene that oxidizes TMA to TMAO. However, it is possible that the regulatory gene affects TMAO concentration in a different manner, such as via the gut microbiome or renal excretion. We also speculated that the variant affects a protein-coding gene, but we cannot rule out other forms of regulation, potentially through miRNA or epigenetic mechanisms related to choline.

## Conclusions

Fasting plasma TMAO concentration is influenced by genetics, sex, and diet, as demonstrated by the QTL on chromosome 12. These factors interact to influence TMAO concentrations, underscoring the importance of considering sex as a biological variable and incorporating environmental exposures, such as diet, into analyses.

## Materials and Methods

### DO Primary-Level Meta-Analysis

We report on the genetic regulation of plasma TMAO concentration using individual-level data from 1,482 female and male mice from five distinct DO studies. Designs of each study are described in detail below. Briefly, for all studies mice were provided a purified diet of AIN-76A (AIN76) at the time-point studied for the meta-analysis and all mice were maintained on a 12-hour light -dark cycle under standard vivarium temperature/humidity with ad lib access to food and water. All DO mice were purchased from Jackson Laboratory (Bar Harbor, ME, USA; J:DO mouse strain, JAX stock #009376) and all study protocols were approved by the university’s Institutional Animal Care and Use Committee (IACUC, University of North Carolina Chapel Hill or the University of California Davis). For all studies, blood was collected from the retroorbital plexus under Isoflurane anesthesia after a 4 hour fast, processed to plasma, and stored at -80°C.

#### Meta-analysis study 1 (DO-UNC1)

DO female and male mice (n=193, female=87, male=106, DO generation=10) were singly housed and fed AIN-76A (D10001, Research Diets, New Brunswick, NJ, USA) throughout the 5-week duration of the study, beginning at eight weeks of age. Blood was collected from unfasted mice.

#### Meta-analysis study 2 (DO-UNC2)

A thorough description of the study design has been published (14,26–28). Relevant to the present meta-analysis, 280 mice were caged in groups of five by sex and fed an AIN-76A for two weeks. Mice were approximately six weeks old at the time of the blood draw and were from DO generation 11.

#### Meta-analysis study 3 (DO-Davis)

DO female mice (n=190) were singly housed in divided cages and fed AIN-76A for two weeks. The study was conducted in two phases including mice from DO generation 26 and 28. At the start of the study, mice from generation 26 were approximately 44 weeks old, and mice from generation 28 were approximately 40 weeks old. Prior to the study, the mice were used as breeders to generate the DO-F1 progeny (see DO-F1 below).

#### Meta-analysis study4 (DO-F1)

A thorough description of the study design has been published (29). Transgenic *CETP/ApoE3* Leiden male mice were bred with DO mice to create 454 female and male *CETP/ApoE3* Leiden x DO F1 (DO-F1) mice. The female DO mice used for breeding were from generations 26 and 28 (DO-Davis, Methods). DO-F1 mice were genotyped to confirm the presence of the *CETP* and *ApoE3* Leiden transgenes. Upon weaning, the DO-F1 mice were provided AIN-76A until eight weeks of age.

#### Meta-analysis study 5 (DO-Choline)

DO mice were purchased in five batches of approximately 40 males and 40 females that spanned 3 distinct DO generations. 155 from generation 42, 142 from generation 43, and 87 from generation 44. Mice were co-housed unless displaying aggression. Mice were provided AIN-76A for two weeks (baseline time point) and were then randomized by cage to receive either an AIN-76A diet supplemented with 1.1% choline chloride (HC, D21042301 Research Diets, New Brunswick, NJ, USA) or to continue the standardized AIN-76A diet for two additional weeks (final time point). The composition of the diets is provided in **Table S1**. Plasma was collected before and after diet randomization. At the final timepoint following blood collection, the mice were euthanized by cervical dislocation under isoflurane anesthesia and tissues were harvested and snap frozen in liquid nitrogen. Snap frozen tissues were stored at -80°C until processed. All animal protocols were approved by the University of California Davis IACUC (Protocol number 21225).

### Genotyping

In all five DO studies, genotyping was performed on DNA isolated from mouse tails submitted to Neogen (Lansing, MI, USA). Genotypes for DO-UNC1 and DO-UNC2 were determined using the Mega Mouse Universal Genotyping Array (MegaMUGA), which contains approximately 70,000 SNP markers. Genotypes for DO-Davis, DO-F1, and DO-Choline were determined using the Giga Mouse Universal Genotyping Array (GigaMUGA), which contains over 143,259 SNP markers designed to capture the diverse genotypes of the Diversity Outbred founding strains (30). In all cases, genotyping was performed on Illumina’s Infinium platform. Genotype calls were converted from nucleotides to A, H, and B, such that A represents major allele homozygous, H represents heterozygous, and B represents minor allele homozygous based on the known genotypes of the founding strains.

For the meta-analysis, which utilized studies with different genotyping depths, a modified approach was used. Each study was evaluated individually to ensure the percentage of genotyped markers per sample was over 90%. Samples were removed if they had over 10% missing markers in their independent study. Due to combining data from the MegaMuga and GigaMuga arrays, genotype probability was calculated with a probability of 0.02%.

### DO-Progenitors

The TMAO values and hepatic gene expression from the DO-Progenitor study has been previously reported (31,32). Briefly 8 female mice from each of the eight mouse strains used to generate the DO model were purchased from the Jackson Laboratory and fed AIN-93M for four weeks. Subsequently 4 of the 8 mice per strain were randomized to remain on the AIN-93M diet (D10012M, Research Diets, New Brunswick, NJ, USA) or fed a high-fat high-cholic acid diet (D12109C, Research Diets, New Brunswick, NJ, USA) for 16 weeks. Tissue, including liver, was collected upon termination via cervical dislocation after anesthesia with isoflurane. Hepatic library preparation, sequencing, mapping, and quantification have been previously described (31). For the candidate gene interrogation, we selected genes with gene expression within ± 1 Mb of the SNP with the highest LOD score in the chromosome 12 QTL identified in this meta-analysis. We tested for differential expression by strain using the Kruskal-Wallis test with Benjamini-Hochberg (BH) corrections and performed pairwise comparisons using the pairwise Wilcox test with BH corrections.

### CAST X BL6 F2 Mouse Model

The development of the CAST X BL6 F2 model has been described (18). CAST and BL6 mice were intercrossed to generate 442 F2 progeny (female=276). Mice were provided standard chow until ten weeks of age, followed by a Western diet (Teklad, 88137) for eight weeks. Mice were fasted overnight before sacrifice and tissues were flash frozen in liquid nitrogen and stored at -80°C until processing. RNA expression was measured on a custom array by Agilent Technologies and consisted of 39,280 oligonucleotides. DNA was isolated using DNeasy tissue kits from QIAGEN (Qiagen, Hilden, Germany) and was performed by Rosetta Inpharmatics (Merck & Co, Rahway, NJ, USA). Genome-wide expression-QTL mapping was performed using the non-parametric Kruskal-Wallis test and corrected for multiple testing using the false discovery rate method. The tests were performed using the NAG C library (33). Empirical permutation tests were used to determine sufficient convergence to estimate significance.

#### ***C***andidate (Acyp1) gene studies

Acyp1 under- and over-expression experiments were performed to test the relationship of the candidate gene and TMAO. Adeno-associated virus 8 vector (AAV8) with a human thyroid hormone binding globulin promoter to target liver-specific transgene expression was purchased from Vector Biolabs (Malvern, PA USA) to over express *Acyp1*. Twenty 4-week-old C57BL/6J mice (10 male and 10 female) purchased from Jackson Laboratory and were allowed to acclimate for 3 weeks then placed on AIN-76A diet. At 10 weeks of age, 5 mice per sex received either a fluorescent control (AAV8-TBG-EGFP) or AAV8-TBG-Acyp1 construct. Injections were given intraperitoneally at a load of 3×10^11^ vg. Blood and liver tissue were collected 2 weeks after the injections and immediately processed to plasma and/or stored at -80°C. TMAO was quantified from plasma as described below and expression of *Acyp1* and *Fmo3* was assessed in liver. RNA was isolated from liver tissue using Qiagen RNeasy kits (Qiagen, Hilden, Germany) and quantified using the Thermo Scientific™ NanoDrop™ One Microvolume UV-Vis Spectrophotometer (ThermoFisher Scientific). Pooled cDNA was used to make a serial standard curve to assess relative sample concentrations corrected by GAPDH for each target. The Powerup SybrGreen method with targeted primers was used to analyze samples on the Applied Biosystems QuantStudio 7 Flex Real-Time PCR system (ThermoFisher Scientific).

*Acyp1* knockout mice were derived from cryopreserved sperm from C57BL/6NJ-Acyp1em1(IMPC)J/Mmjax, RRID:MMRRC_066809-JAX from the Mutant Mouse Resource and Research Center (MMRRC) at The Jackson Laboratory, an NIH-funded strain repository, and was donated to the MMRRC by Stephen Murray, Ph.D., The Jackson Laboratory. Twenty-two mice (11 male and 11 female; 12 *Acyp1*^-/-^ and 10 *Acyp1*^+/+^) were fed AIN-76A for two weeks and then plasma and tissues were collected. RNA expression of *Acyp1* and *Fmo3* was assessed similarly to the over-expression experiments. TMAO was quantified from plasma as described below.

### Hepatic Gene Expression

Candidate gene expression was measured on hepatic RNA from male and female mice from the DO-Choline study. Liver RNA was isolated using the Applied Biosystems MagMAX mirVana Total RNA Isolation Kit (ThermoFisher Scientific, Waltham, MA) and KingFisher DuoPrime Extraction System (ThermoFisher Scientific). RNA concentration and purity were checked by the Thermo Scientific™ NanoDrop™ One Microvolume UV-Vis Spectrophotometer (ThermoFisher Scientific) and the Agilent 2100 Bioanalyzer (Agilent, Santa Clara, CA). Isolated RNA was diluted to 50 ng/μL with nuclease-free water and synthesized into cDNA using the Applied Biosystems High-Capacity cDNA Reverse Transcription kit (ThermoFisher Scientific). 3 μL of each cDNA sample was pooled, and a standard curve was generated from serial dilutions.

Quantitative polymerase chain reaction (qPCR) was conducted with PowerUp™ SYBR™ Green reagent and custom primers specific to each target gene. Primers were diluted from 100 μM to 10 μM prior to use. The qPCR master mix for each sample well consisted of 0.5 μL of forward primers, 0.5 μL of reverse primers, 2.5 μL of nuclease-free water, and 5 μL of PowerUp™ SYBR™ Green. The Eppendorf Epmotion 5075 Liquid Handler robotic system (Eppendorf, Hamburg, Germany) was used to add qPCR master mix, standards, and samples to the 384-well plate in singlet. The 384-well qPCR plate was run on the Applied Biosystems QuantStudio 7 Flex Real-Time PCR system (ThermoFisher Scientific). Each gene was run three times on three qPCR plates to have quantities calculated in triplicate. ACYP1 abundance was calculated from the pooled cDNA standard curve and normalized to GAPDH and expressed as fold of control.

### TMAO Quantification

TMAO was measured by liquid chromatography mass spectrometry using a method adapted from Wang and colleagues (4). Deuterated analytes were used to generate 10 µM surrogate standards (SSTD) that were spiked into each sample, standard, and control. A standard curve ranging from 100 µM to 0.0475 µM was used. A human plasma sample was aliquoted and run as a control. Additionally, a sample at the beginning, middle, and end of each LCMS run was extracted in duplicate and served as an extraction control. To prepare the samples, plasma from fasted mice was thawed on ice and 20 µL of plasma was combined with 80 µL of 10 µM SSTD and vortexed for 30 seconds. Samples were centrifuged for 10 minutes at 10°C at 18,000 g. The supernatant was transferred filtered using Ultrafree MC Centrifugal filter tubes (Durapore PVDF 0.1µM) at 12,000 g for 4 minutes at 10 C then transferred to HPLC vials with 150 µL inserts. Metabolites were measured using liquid chromatography mass spectrometry on an Acquity UPLC and API 4000 Q-Trap with a 150 x 2 mm silica column containing 3 µm particles (Phenomenex, Torrance, CA, USA). A discontinuous gradient of water:0.1% acetic acid (solvent A) and methanol:0.1 acetic acid (solvent B) was employed over ten minutes. For each sample, a ratio of 98% A:2% B was increased linearly to 85% A:15% B over five minutes. From minute 5 to 6:15, the ratio changed to 100 % solvent B, which was held until minute 8. The ratio then returned to 98%A:2%B until minute 10.

### Statistical Analysis

#### Quantitative Trait Locus Analysis

Quantitative trait locus analysis was completed using the library qtl2 in RStudio (qtl2 version 0.24) (39). A hidden Markov model with transition states was used to determine genome probabilities. Genome probabilities were collapsed from 36 diplotype states (AB, AC, AD. HH) to 8 allele probability states (A-H). A linear mixed model (lmm) including a kinship matrix generated using the “Leave-One-Chromosome-Out” (kLOCO) method was used (40). In the DO-Choline study, models including male and female mice were adjusted for sex and experimental batch, whereas models with only male or female mice were only adjusted for experimental batch. In the meta-analysis, models were adjusted by study instead of experimental batch. All covariates were numerically coded and covariates with more than two levels were coded using matrix multiplication. In DO-Choline at the final time point, the diet arms were assessed individually by including the mice randomized to a given arm. Phenotypes were transformed to comply with the normal distribution using the natural log transformation or the square root. A logarithm of odds (LOD) score was calculated by comparing a lmm including genotype, covariates, and kinship to a lmm excluding genotype but including covariates and kinship. To determine significant SNP-phenotype relationships, 1000 permutations were performed. For a phenotype, one permutation consisted of randomizing the genotypes, calculating a LOD score for each SNP, and recording the highest genome-wide LOD score observed by chance. After repeating this process 1000 times, the 95% Bayesian confidence interval was calculated to determine the LOD significance threshold for that phenotype. QTL confidence intervals were also defined by the 95% Bayesian credible interval.

#### Heritability

Narrow sense heritability was calculated using the est_herit() function in R/qtl2. In the meta-analysis and in studies with both sexes, sex and a study-specific square kinship matrix were used as covariates. In studies with only females (DO-UNC2, DO-Davis) only kinship was included.

#### Assessments of Peak Marker

To determine the genotypes or strain-related regression coefficients at the peak genetic marker in a QTL, the marker was first identified using the find_marker() function in the R package qtl2. For genotype assessments, the peak marker was identified in the raw genotype file and merged with the phenotype data. For determining study-specific strain-related regression coefficients, the linear model was run for each study independently, and the results were subset to the peak marker identified by find_marker().

#### Allele Distribution

To determine the relative distribution of the founding alleles per chromosome, allele probabilities calculated via qtl2 were used. For each marker on a given chromosome, the sum of the allele probabilities was calculated (A - H). This process was repeated for each marker on a given chromosome. Then, the values were summed per allele and divided by the total allele probability scores to get a relative percent. This process was repeated for each chromosome.

#### Statistics

Phenotypes were assessed for compliance with the normal distribution using the Shapiro-Wilk test. Phenotypes were considered compliant if they produced a W-statistic greater or equal to 0.95. Transformations were considered in the order of natural log, square root, and rank. TMAO was natural log transformed for QTL analyses. Differences in the phenotypic mean were tested using ANOVA followed by Tukey’s post-hoc test.

To test for the sex by genotype interaction, the marker with the highest LOD score in the QTL was selected. An ANOVA model assessing the interaction of sex and genotypes was tested with the genotypes coded as factors (A = homozygous for the major allele, H = heterozygous, B = homozygous for the minor allele). To complete post-hoc testing, the R package emmeans (v 1.8.1-1) was used with the method “specs” sex and genotype.

## Acknowledgements

We would like to acknowledge the contributions from the UC Davis Meyer Hall Vivarium including Cheryl Perez and Sue Bennett for their assistance with the DO-Choline project. We would like to acknowledge the help of Anita Wen for participating in the DO-Choline dissections and Korrie Tugal for her work as a volunteer laboratory assistant. We would also like to acknowledge Karl Broman for providing feedback on key steps of the meta-analysis via the R/qtl2-discussion Google Group.

## Financial Disclosure Statement

This work was funded by the USDA, Agricultural Research Service (2032-51530-025-00D), and the National Institute of Health (R01HL148110, R01HL168493, U54HL17032, and R01DK143650). The funders had no role in the study design, data collection and analysis, decision to publish, or preparation of the manuscript.

## Competing Interests

The authors declare that the research was conducted in the absence of any commercial or financial relationships that could be construed as potential conflicts of interest.

## Data Availability Statement

The data underlying this article are available in Dryad Digital Repository, at https://dx.doi.org/[doi] (made accessible after acceptance).

## Notes

### Competing Interest Statement

The authors have declared no competing interest.

